# VviMYB41 orthologs contribute to the water deficit induced suberization of grapevine fine roots

**DOI:** 10.1101/2020.05.06.080903

**Authors:** Li Zhang, Isabelle Merlin, Stéphanie Pascal, Pierre-François Bert, Frédéric Domergue, Gregory A. Gambetta

## Abstract

The permeability of roots to water and nutrients is controlled through a variety of mechanisms and one of the most conspicuous is the presence of structures such as the Casparian strips and suberin lamellae. Roots actively regulate the creation of these structures developmentally, along the length of the root, and in response to the environment, including abiotic stresses such as drought. In the current study, we characterized the suberin composition along the length of grapevine fine roots during development and in response to water deficit. In parallel samples we quantified changes in expression of suberin biosynthesis- and deposition-related gene families (via RNAseq) allowing the identification of drought-responsive suberin-related genes. Grapevine suberin composition did not differ between primary and lateral roots, and was similar to that of other species. Under water deficit there was a global upregulation of suberin biosynthesis which resulted in an increase of suberin specific monomers, but without changes in their relative abundances, and this upregulation took place across all the developmental stages of fine roots. These changes corresponded to the upregulation of numerous suberin biosynthesis- and deposition-related genes which included orthologs of the previously characterized AtMYB41 transcriptional factor. Functional validation of two grapevine MYB41 orthologs, VviMYB41 and VviMYB41-like, confirmed their ability to globally upregulate suberin biosynthesis and deposition. This study provides a detailed characterization of the developmental and water deficit induced suberization of grapevine fine roots and identifies important orthologs responsible for suberin biosynthesis, deposition, and its regulation in grape.

**One sentence summary:** Our study details the biochemical changes and molecular regulation of how grapevines decrease their root permeability during drought.

## INTRODUCTION

The root system is highly dynamic in space and time, and roots actively regulate water and nutrient uptake during development and in response to the environment. This regulation occurs across different time frames. There are rapid changes over the short-term (*i.e.* hours to days), for example the modulation of root water uptake via aquaporins (Gambetta et al., 2017), or nutrient uptake via transporters (Williams and Miller, 2001), or slower more long-term changes (*i.e.* days to months to seasons) such as the modification of root system architecture (Osmont et al., 2007) and structural changes in individual roots.

Within the individual root, one of the most well-established structural features impacting water and nutrient uptake is the development of the Casparian strips and the suberin lamellae in the endodermis and exodermis of primary roots. Only the root tip (typically just the first few mm) lacks any suberized structures, so the Casparian band and suberized endo/exodermis develop relatively rapidly, and in woody perennials this development continues with the formation of a woody, highly suberized periderm (Evert, 2006). Therefore, as these suberized structures become more developed in root portions further away from the root tip (*i.e.* older parts of the roots) they contribute to decrease the hydraulic conductivity (Kramer and Bullock, 1966; MacFall et al., 1991; Gambetta et al., 2013) and the passive uptake of nutrients (Ranathunge et al., 2011). In response to environmental factors like drought, water logging, and salt stress there is an increased deposition of suberin lamellae which contributes to further decreases in root permeability (Franke and Schreiber, 2007; Ranathunge et al., 2011; Aroca et al., 2012).

Suberin is a very complex heteropolymer made of polyphenolic and polyaliphatic domains together with non-covalently associated waxes (Bernards, 2002; Delude et al., 2016). Suberin is deposited at the interface between the plasma membrane and the primary cell wall. Although the fatty acid-derived monomers making the aliphatic network and waxes have now been characterized in several plants and tissues, the macromolecular structure of the polymer, the domain interconnections, as well as the interactions of suberized layers with cell wall constituents remains elusive. Visualized with transmission electron microscopy, suberin often appears as a lamellar structure made of 5 to 10nm thick electron-translucent and -dense alternating bands. Latest models based on ^13^C ssNMR studies suggest that light bands correspond to membrane-like ordered aliphatic-glycerol esters, while dark bands are made of hydroxycinnamic acids, especially ferulic acid, which are extensively covalently-linked in a lignin-like macromolecule (Graça, 2015). Measuring the permeability of suberized cell walls present in roots using transport chambers would require the enzymatic isolation of the suberized barriers present in the endodermis and/or exodermis, which is currently not technically possible (Schreiber, 2010). As an alternative, most studies on suberin permeability were conducted using potato (*Solanum tuberosum*) tuber skin, which is composed of about ten compacted layers of suberized cells. It was shown that the water permeability was inversely correlated with the post-harvest time because of the significant amounts of waxes that were deposited upon storage (Schreiber et al., 2005b). These analyses suggested that in aerial cuticles, the barrier properties of potato skin were more related to the deposition of waxes than to the lipophilic polymer. Although technically challenging, the radial transport of water and solutes has been assessed in rice and corn primary roots using pressure probes (Ranathunge et al., 2005; Schreiber et al., 2005a; Schreiber et al., 2005b). These approaches indicated that the root suberized cell walls are at least 3 orders of magnitude more permeable than potato skin or plant cuticles. This much higher permeability relative to aerial cuticles could be related to the fact that in roots suberized layers are not impregnated with high amounts of waxes. This is also congruent with the idea that the primary role of roots is water uptake, thus highly (or perhaps perfectly) impermeable suberized walls in roots should be deleterious to plants.

In the past, studies on suberin were restricted to the periderms of cork oak tree and potato tubers, but in the early 2000s Arabidopsis, and more recently agonomically important fruits like tomato (Lashbrooke et al., 2015), apple (Legay et al., 2017), and watermelon (Cohen et al., 2019) became models of choice for large genomic studies. Gas-chromatography based analysis methods showed that the polyaliphatic domain of suberin is principally made of a complex mixture of α,ω-bifunctional fatty acids (principally ω-hydroxyacids and α,ω-diacids) and glycerol which are primarily linked by ester linkages into a polyester. Various proportions of α,ω-bifunctional fatty acids differing in term of chain length, degrees of unsaturation, and oxidation generate an aliphatic suberin composition which is often specific to each plant, and even to the different suberized layers of the same plants. Together with various transcriptomic studies conducted on suberizing tissues candidate biosynthetic genes were identified for suberin aliphatics, allowing a metabolic pathway to be proposed (reviewed in Vishwanath et al., 2015). In this hypothetical biosynthetic pathway, 18-carbon atoms fatty acid precursors are elongated in the endoplasmic reticulum by elongase complexes, in which the keto-acyl-CoA synthase (KCS) plays the major role by controlling the final chain-length (Franke et al., 2009). Fatty acids ranging from 16 to 24 carbon atoms are oxidized by cytochrome P450 (CYP86A1 and B1) to omega-hydroxy fatty acids (Höfer et al., 2008; Compagnon et al., 2009), and eventually to dicarboxylic acids, this last step remaining uncharacterized to date. Oxidized aliphatics are then transferred to glycerol by *sn-2* specific glycerol-3-phosphate acyl-transferase (GPAT; Beisson et al., 2007), yielding the putative building blocks of the polymer. These acyl monomers (or oligomers) are then exported outside of the cell through putative ATP binding cassette transporter of the G-clade (ABCG; Yadav et al., 2014). Long-chain acyl-CoA synthase (LACS) are apparently also important for suberin aliphatic synthesis even if their exact role and position within this pathway remain elusive (Pollard et al., 2008). Additionally, BAHD-acyl transferase like aliphatic suberin feruloyl transferase (ASFT) are important for the incorporation of hydroxycinnamic acids in the polymer (Molina et al., 2009). Finally, at the level of transcriptional regulation several MYB transcription factors (MYB9, MYB41, MYB93 and MYB107) have been shown to be important positive regulators of suberin biosynthesis (Kosma et al., 2014; Lashbrooke et al., 2016; Legay et al., 2016; Gou et al., 2017; Cohen et al., 2020).

In the current study, we characterized the suberin composition along the length of grapevine fine roots examining four distinct root portions, each representing a specific developmental stage, in order to quantify changes in suberin composition during fine root development. These analyses were paired with roots that had been subjected to two different levels of water deficit. We characterized the orthologous gene families of known suberin biosynthesis- and deposition-related gene families in grape and examined their expression (via RNAseq) in parallel samples of fine roots subjected to water deficit. This allowed the identification of grape orthologs of the suberin biosynthetic regulator MYB41 transcription factor whose expression was highly upregulated under water deficit. Finally, the functionality of these *Vitis* orthologs was confirmed via transient expression in agroinfiltrated *Nicotiana benthamiana* leaves.

## RESULTS

### Suberin Composition of Grapevine Primary and Lateral Roots

Grapevine fine roots were exhaustively delipidated, the suberin polyester depolymerized by acidic transmethylation, and its monomer composition and content analyzed by gas chromatography. The polyester composition was dominated by typical suberin monomers such as dicarboxylic acids (DCA) and omega hydroxy acids (ωOH) with chain lengths ranging from C16 to C22 (Fig. 1A and B). In both primary and lateral roots, four major monomers (16:0-DCA, 18:1-DCA, 18:1-ωOH and 22:0-ωOH) represented more than 50% of the total content (Fig. 1A), and dicarboxylic and omega hydroxyl acids collectively accounted for about 70% of the total (Fig. 1B). Other monomers detected were fatty acids, fatty alcohols, and diols. Interestingly, diols (especially C18:0- and C20:0-diol) were more abundant in the primary root than in lateral roots (12.7 and 4.9% of the total, respectively). Concerning the acyl chain lengths, 18-carbon molecules represented more than 40% of the total suberin acyl-chains in both types of roots, while molecules with 16, 20, and 22 carbon atoms each accounted for about 20% of the total (Fig. 1C). Although primary and lateral roots had a rather similar acyl-chain composition, the total monomer content in lateral roots was 20% lower (12.9 µg/mg DR in the primary root *vs* 10.3 µg/mg DR in lateral roots).

**Figure 1.**
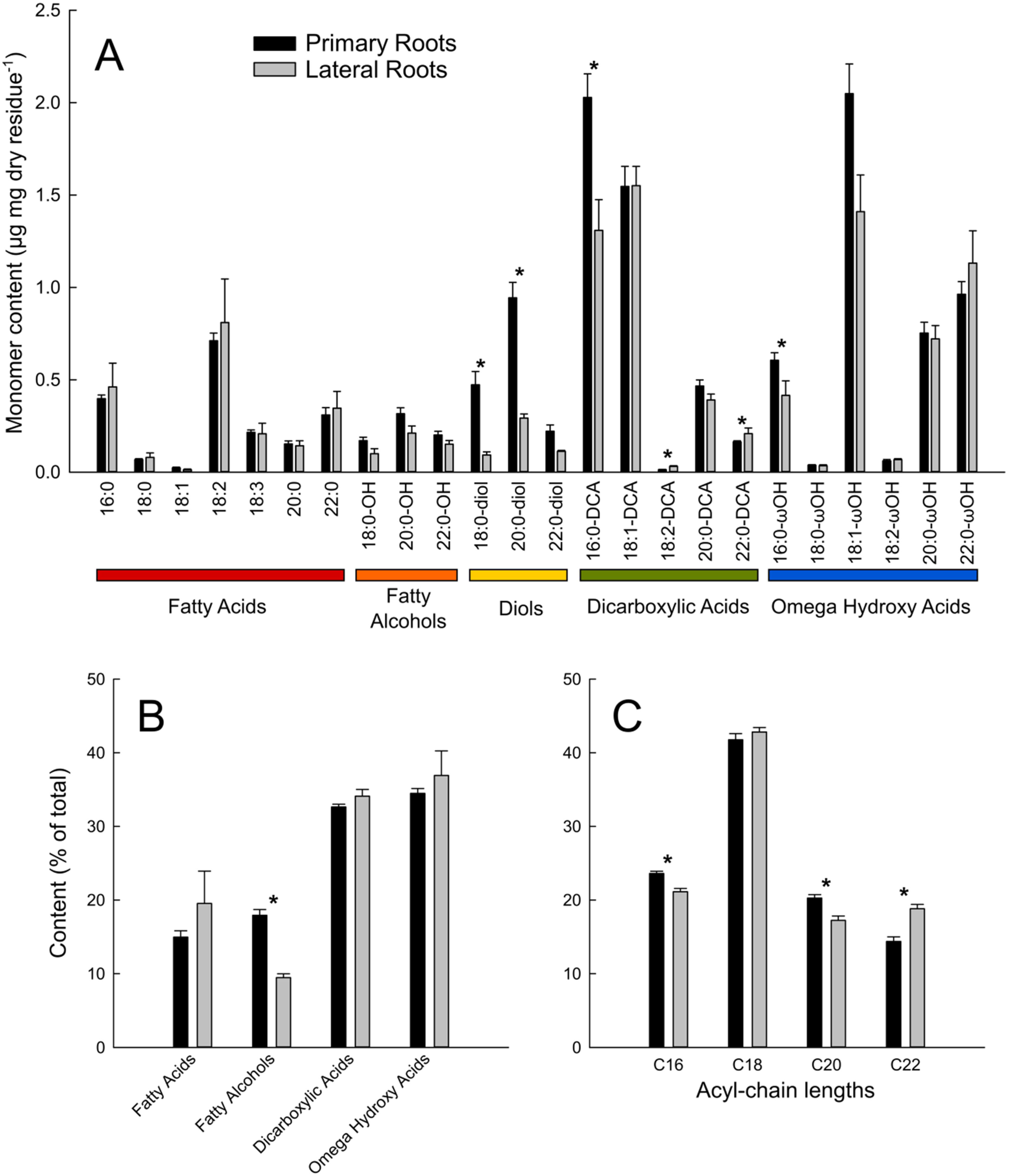
Aliphatic suberin composition of well-watered primary (black bars) and lateral (grey bars) grapevine roots. Suberin monomers from solvent-extracted roots were released by acidic transmethylation, silylated, separated by gas chromatography and quantified using internal standards. A, Suberin monomer composition in µg/mg dry residue. B, Types of acyl-chains and C, Acyl-chain length distributions (in % of total). Error bars represent ± standard error and different letters designate statistically significant differences (n=10 primary roots and n=3 lateral roots; P<0.05 TUKEY’s HSD).

### Developmental Changes in Suberin Composition

We analyzed the global lipid composition of four different developmental stages of the primary root: the root tip, and segments corresponding to the development of Primary Xylem, Early Secondary Growth (prior to periderm development), and Late Secondary Growth (which included a periderm), where the developmental stage of each analyzed sample was confirmed by microscopy. Global lipid analysis allowed for the quantification of the totality of the acyl-chains present in the various root segments, including the major suberin monomers, as well as that of sterols and hydroxycinnamates such as coumaric, ferulic, and synapic acids (Fig. 2A). Among the different acyl-chains detected, fatty acids mainly come from membrane lipids, 2-hydroxy acids are a signature from the sphingolipids present in the plasmalemma, while dicarboxylic acids, omega hydroxy acids, fatty alcohols, and diols represent suberin-specific aliphatics. It should be pointed out that this global lipid analysis slightly underestimates the total suberin monomer content as the solvent-extraction method indicated the presence of classical fatty acids in the suberin polyester (Fig. 1).

**Figure 2.**
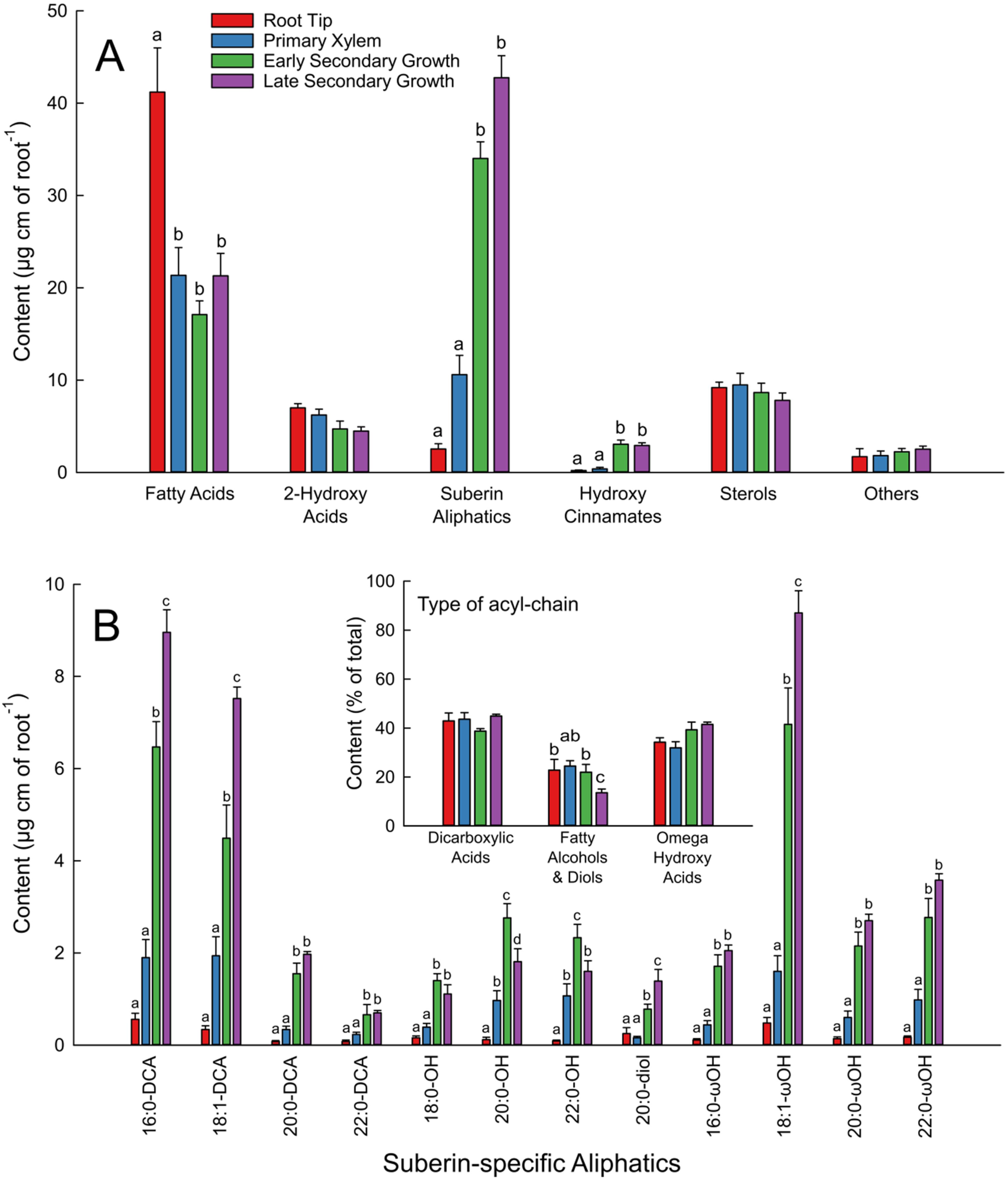
Global lipid analysis of different portions of well-watered primary roots coorsponding to the root tip (red bars), and the developmental stages of primary xylem (blue bars), early seconday growth (green bars), and late secondary growth (purple bars). Root portions were directly transmethylated, silylated, and the major lipid components were separated by gas chromatography and quantified using internal standards. A, Global lipid composition in µg/linear cm of each root portion. B, Suberin-specific aliphatic compostion in µg/linear cm of roots. B inset, Distribution of suberin acyl-chain types (in % of total). Error bars represent ± standard error and different letters designate statistically significant differences (n=5-6; P<0.05 TUKEY’s HSD).

Fatty acids predominated in the root tip (~40 µg per linear cm of root) and were about the same (~20 µg per linear cm of root) in the Primary Xylem, and Early and Late Secondary Growth stages (Fig. 2A). The root tip contained very few suberin-specific aliphatics (2.5 µg per linear cm of root) and high amounts of fatty acids (42.6 µg per linear cm of root). Considering all the suberin aliphatic classes together, their content increased dramatically with the age of the root segment: from 10.6 µg per linear cm of root in the Primary Xylem zone up to 59.8 µg per linear cm of root in the and Late Secondary Growth with periderm zone (Fig. 2A and B). In contrast, the amounts of most 2-hydroxy acids and sterols did not change across the different developmental stages of the root (Fig. 2A; Supplemental Fig. S1). Although the total suberin monomer content was significantly different in the various developmental stages, its composition displayed much less variation. Dicarboxylic acids systematically represented about 39 to 45% of the total monomers, omega hydroxy acids 32 to 41% while fatty alcohols and diols 14 to 24% (Fig. 2B, inset). Detailed quantification of unsubstituted fatty acids, hydroxycinnamates, 2-hydroxy fatty acids, and sterols is presented in Supplemental Figure 1.

### Changes in Suberin Composition in Response to Water Deficit

We subjected grapevines to different watering regimes (control, medium, and high water stress). Water deficit decreased vine water status with average pre-dawn water (Ψ_Predawn_) potentials decreasing from near zero in controls to approximately −0.5 and −1.5MPa under medium and high water deficit respectively (Fig. 3A). Under medium water deficit there was an approximate 80% decrease in stomatal conductance and 60% decrease in assimilation, both of which decreased to nearly zero under high water deficit (Fig. 3B and C). In addition, root hydraulic conductivity (Lp_root_) was assessed. Water deficit resulted in decreases in Lp_root_ of approximately 50% under medium deficit and 80% under high deficit (Fig. 3D). Fine roots were stained with the fluorescent dye berberine hemisulfate in order to visualize suberized structures in the roots (Fig. 4). Water stressed roots developed a Casparian band in closer proximity to the root tip (Fig. 4A and D) and generally exhibited a greater amount of berberine hemisulfate fluorescence in the endodermis and exodermis.

**Figure 3.**
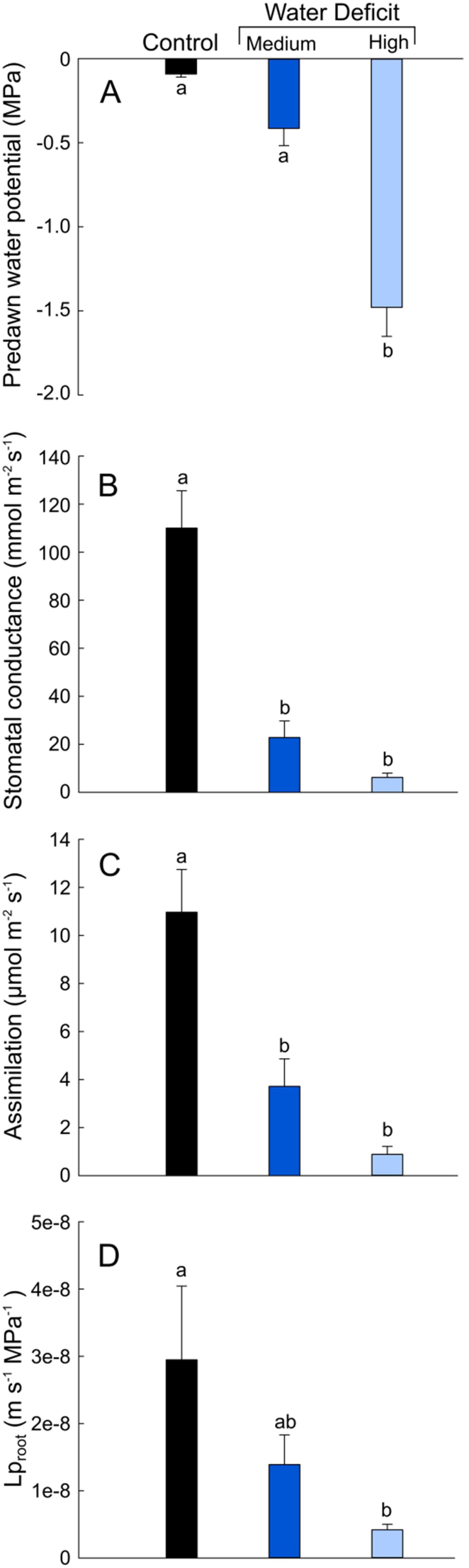
Physiological parameters of grapevines subjected to water deficit. A, Average pre-dawn water potentials; B, stomatal conductance; C, carbon assimilation; and D, root hydraulic conductivity (Lp_root_) for plants that were well-watered (black bar), or subjected to medium (royal blue bar) or high (light blue bar) levels of water deficit. Error bars represent ± standard error and different letters designate statistically significant differences (n=5-7; P<0.05 TUKEY’s HSD).

**Figure 4.**
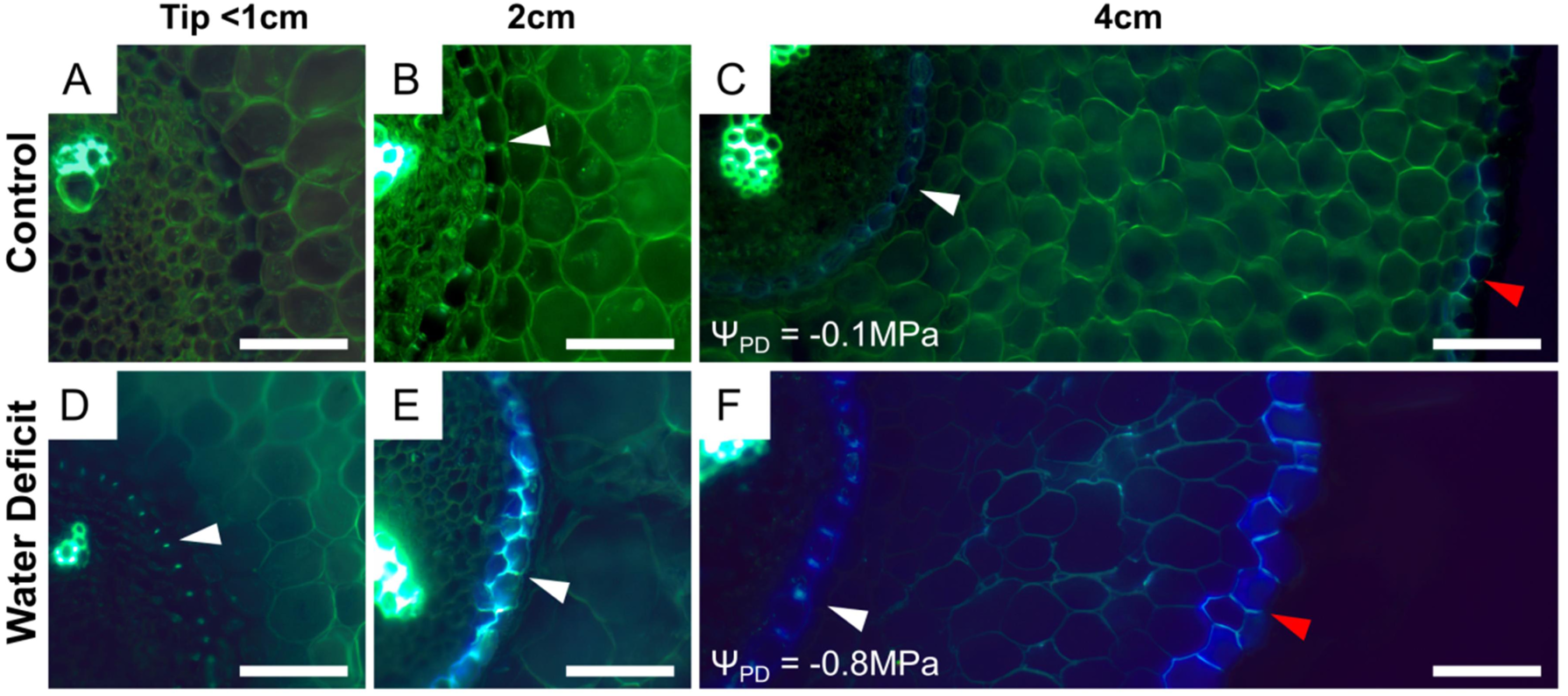
Berberine hemisulfate staining of root cross sections. A-F, Representative images of RGM fine roots grown under control (A-C) and water deficit (D-F) conditions. Distances from the root apex are noted above each column. The pre-dawn water potentials (Ψ_PD_) of the individual roots shown are given which corresponds to a “Medium” water deficit. White arrows indicate casparian strips in the endodermis (B-F) and red arrows (C, F) indicate the exodermis. White scale bars are 50µm.

We then compared the global lipid composition of roots that had been subjected to the different water regimes across the same four different developmental stages described above. All classical fatty acids as well as 2-hydroxy acids were grouped in order to evaluate the membrane aliphatic content of the different samples. Roots tips were more enriched in membranous aliphatics than the other root segments, which contained about similar levels, and the total amounts of membranous aliphatics increased in all segments with increasing water stress levels (Fig. 5A). Most importantly, the suberin aliphatic content increased with the water stress level in the four different root segments (Fig. 5B). As total suberin was shown to increase with the age of the root segment (Fig. 2), the increases in suberin aliphatic content due to water deficit expressed as a fold-change were much greater in younger root tissues. In the root tip for example, the total amount of suberin aliphatics was respectively two and six times higher in medium and high water stressed roots than in controls. Similarly in the primary xylem developmental stage, suberin increased by about 180-200%. In the older parts of the roots (*i.e.* in zones corresponding to early and late secondary growth) these increases remained highly significant, with approx. 10% and 50% more suberin aliphatics in medium and high water stressed roots than in controls. When looking at the different aliphatic classes comprising the suberin polyester, very similar variations were observed for dicarboxylic and omega-hydroxy acids; the content systematically increased with root age as well as with water status stress level (Fig. 5C and D). In contrast, the fatty alcohol and diol levels showed much less variation (Fig. 5E). Although the amount of these compounds slightly increased in the root tip and primary xylem stages under water deficit, it remained unchanged in older part of the roots. In summary, water stress predominately stimulated the biosynthesis of dicarboxylic and omega-hydroxy acids, while that of fatty alcohols and diols was less, or not, affected.

**Figure 5.**
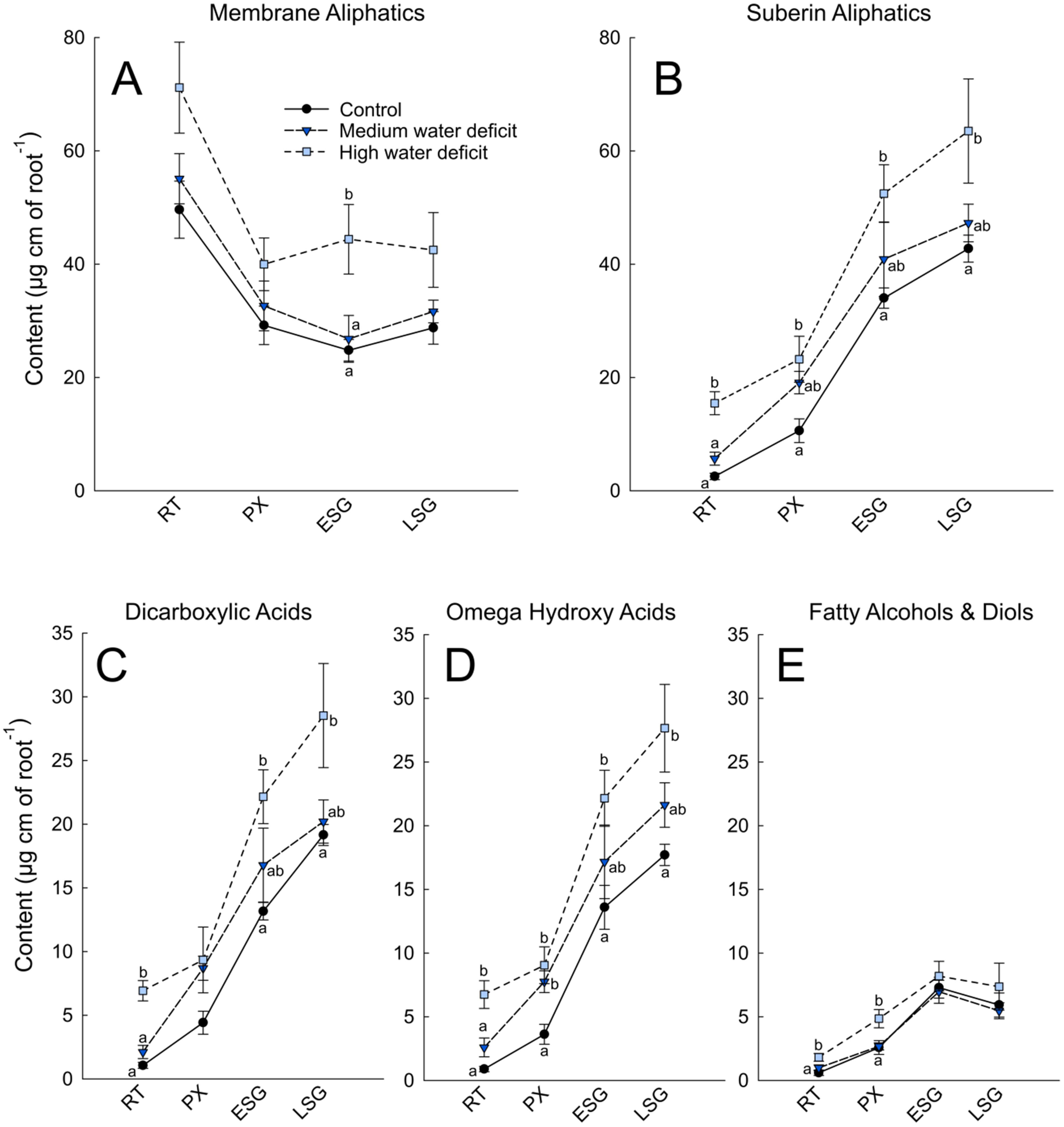
Global lipid analysis of different roots portions coorsponding to the root tip (RT), and the developmental stages of primary xylem (PX), early seconday growth (ESG), and late secondary growth (LSG) under well-watered (black circles), medium water deficit (dark blue triangles) and high water deficit (light blue squares) conditions. Root segments were directly transmethylated, silylated, and the major lipid components were separated by gas chromatography and quantified using internal standards. A, Membrane aliphatics; B, Suberin-specific aliphatics; C, Dicarboxylic acids; D, Omega hydroxy acids; and E, Fatty alcohols and diols content in µg/linear cm of root. Error bars represent ± standard error and different letters designate statistically significant differences (n=4-6; P<0.05 TUKEY’s HSD).

### Transcriptional Regulation of Suberin Biosynthesis-related Gene Families

We utilized the previously characterized genes involved in suberin biosynthesis and deposition (Vishwanath et al., 2015), together with some core wax biosynthesis gene families recently identified and characterized in *Vitis vinifera* (Dimopoulos et al., 2020), to identify orthologous gene families in grapevine. This allowed for a targeted selection of orthologous grapevine genes involved in suberin biosynthesis and deposition (Summarized in Fig. 6; Supplemental Fig. S2). Eighty-one genes were selected across 11 gene families (Supplemental File 1). We carried out RNAseq expression analyses from parallel samples (*i.e.* different roots from the same plants) as those used for the suberin analysis, resulting in RNAseq data for control, medium and high water deficit roots.

**Figure 6.**
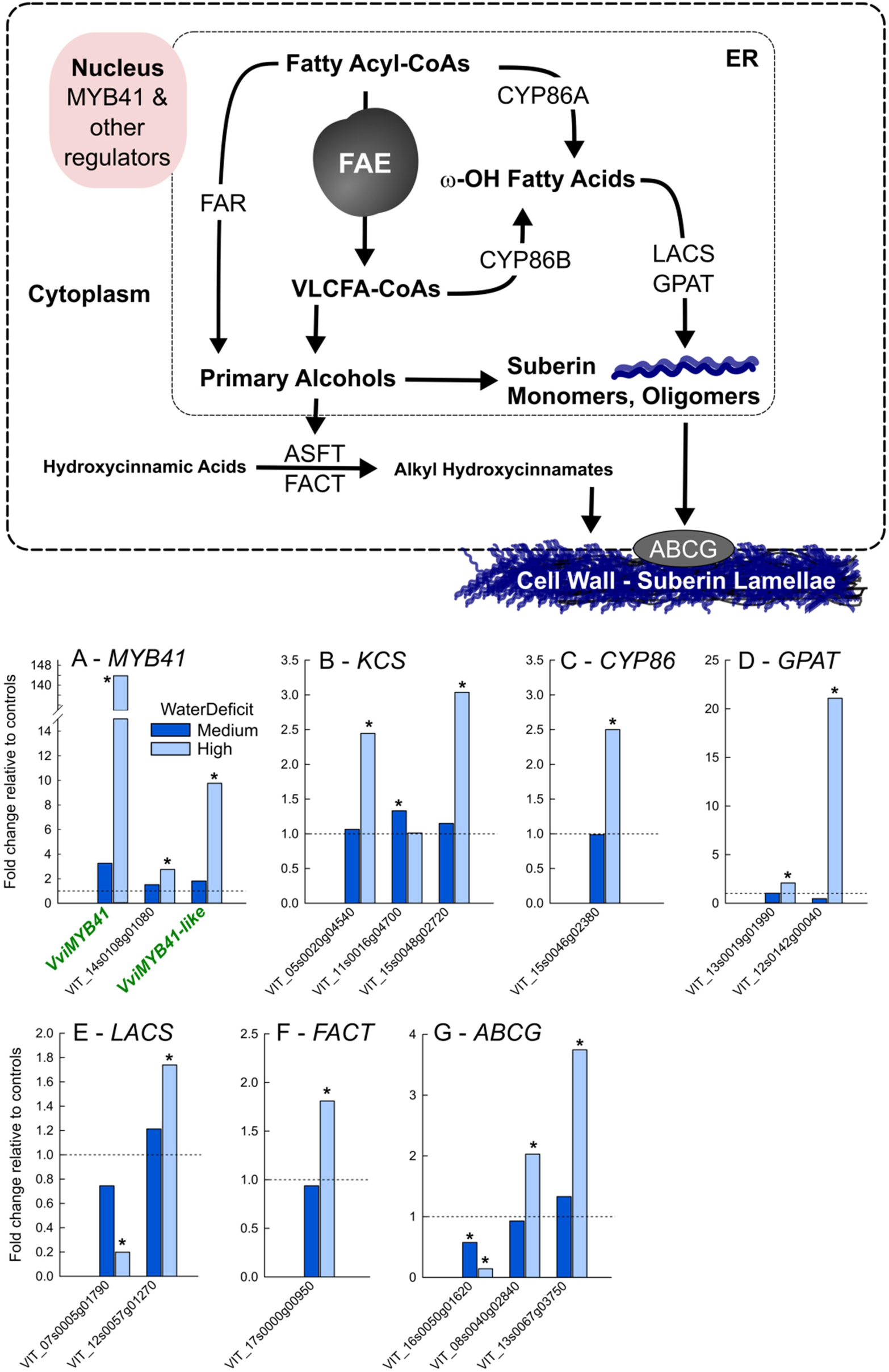
Summary of the changes in gene expression of selected suberin biosynthesis and deposition related genes in grapevine fine roots under water deficit. **Above**, figure (adapted from Vishwanath et al. 2015) summarizing the current knowledge of the gene families involved in suberin biosynthesis and deposition. The gene families assessed in this study included the Myb41 transcription factors, genes comprising the Fatty Acid Elongation (FAE) complex (KCS, KCR, PAS2, and CER10), CYP86s, FAR, LACS, GPAT, ASFT, FACT, and ABCG. **Below**, graphs present the fold change in gene expression in response to medium (green bars) and high (red bars) levels of water deficit relative to controls. Horizontal dotted line represents the expression level of controls. Only gene expression data from genes exhibiting a statistically significant change in expression are shown. Complete FPKM expression data for all genes assessed can be found in Supplemental Fig. S3 and Supplemental File 1. Asterisks designate statistically significant differences relative to controls (n=6 controls and n=3 each water stress level, P<0.05 TUKEY’s HSD).

In total 48 of the 81 genes showed expression in fine roots and certain members of all gene families were represented (Supplemental Fig. S3). Some families only have a single member that was expressed in fine roots, like the FARs (VIT_06s0080g00120) and FACT (VIT_17s0000g00950). Under our experimental conditions, the expression of 15 of these 48 genes (~30%) changed significantly in response to water deficit (Fig. 6; Supplemental Fig. S3). Most of the significant changes in expression involved genes being upregulated under water deficit (13/15, ~85%) and specifically under high water deficit. Notably two grapevine *VviMYBs* that clustered with the Arabidiopsis *AtMYB41* (At4g28110), referred to in this work as *VviMYB41* (VIT_12s0134g00570) and *VviMYB41-like* (VIT_00s0203g00070) based on homology (Supplemental Fig. S2A), were significantly upregulated under water deficit (Fig. 6A). The strongest orthologue, *VviMYB41*, was upregulated more than 100-fold under high water deficit. Of the other *VviMYBs* that clustered with other implicated MYB regulators, *AtMYB39*, *AtMYB92*, *AtMYB107*, and *MdMYB93*, one *Vitis* ortholog, VIT_14s0108g01080, was also significantly upregulated under water deficit (Fig. 6A; Supplemental Fig. S2A). Other notable genes that were upregulated under water deficit included orthologs of *KCS*, *CYP86*, *GPAT*, *LACS*, *FACT*, and *ABCG* genes, which comprise all the major steps of suberin monomer biosynthesis and export (Fig. 6).

### Functional Validation of the VviMYB41 Orthologs

In order to confirm that VviMYB41 and VviMYB41-like transcriptionally regulate suberin deposition, we transiently agroinfiltrated *Nicotiana benthamiana* leaves with vectors overexpressing (35S) *VviMYB41* or *VviMYB41-like*, and 5 days later performed chemical analysis of leaf polymer composition and content (Fig. 7). Both transcription factors led to the accumulation of high amounts of suberin monomers like 18:1 dicarboxylic acid, 22:0 fatty acid, and 22:0 fatty alcohol, which are present in much lower amounts in the cuticle of *Nicotiana benthamiana* leaves as illustrated by the pBIN19 controls (Fig. 7A). *VviMYB41* and *VviMYB41-like* expression also led to the appearance of monomers that are normally absent in leaf cutin, especially the very long chain (VLC) 20:0, 22:0, and 24:0 omega hydroxy and dicarboxylic acids, which are characteristic monomers of suberin. In contrast, both transcription factors had no effect on the typical cutin monomer (10,16) dihydroxy palmitic acid, 16:0diOH (Fig. 7A). While VviMYB41 and VviMYB41-like only slightly increased long chain (C16 and C18) fatty acids, they tremendously increased the amounts of VLC-fatty acids (530 and 1120%, respectively), fatty alcohols (4270 and 3420%, respectively), dicarboxylic acids (760 and 2770%, respectively), and to a lesser extend omega hydroxy acids (180 and 360%, respectively; Fig. 7B). Notably, although both MYBs similarly affected the fatty alcohol content VviMYB41-like had a much stronger effect on the accumulation of omega hydroxy acids, and especially dicarboxylic acids, the amount of 18:1-DCA being increased more than 22 times by *VviMYB41-like* transient expression (Fig. 7A and B). We also analyzed the suberin composition of *Nicotiana benthamiana* well-watered primary roots for comparison (Supplementary Figure S4). When comparing the relative proportions of the five most abundant suberin monomers found in *Nicotiana benthamiana* roots and leaves agroinfiltrated with VvMYB41 or VvMYB41like (Fig. 7C), it appeared that the long chain monomers (16:0DCA, 18:1DCA and 18:1ωOH) were less abundant in agroinfiltrated leaves than in root suberin, whereas the very long chain monomers (22:0 and 22:0ωOH) were enriched in agroinfiltrated leaves when compared to root suberin. This difference may result from the fact that in leaves the epidermis is well-known to have a highly active fatty acid elongation pathway for cuticular wax production.

**Figure 7.**
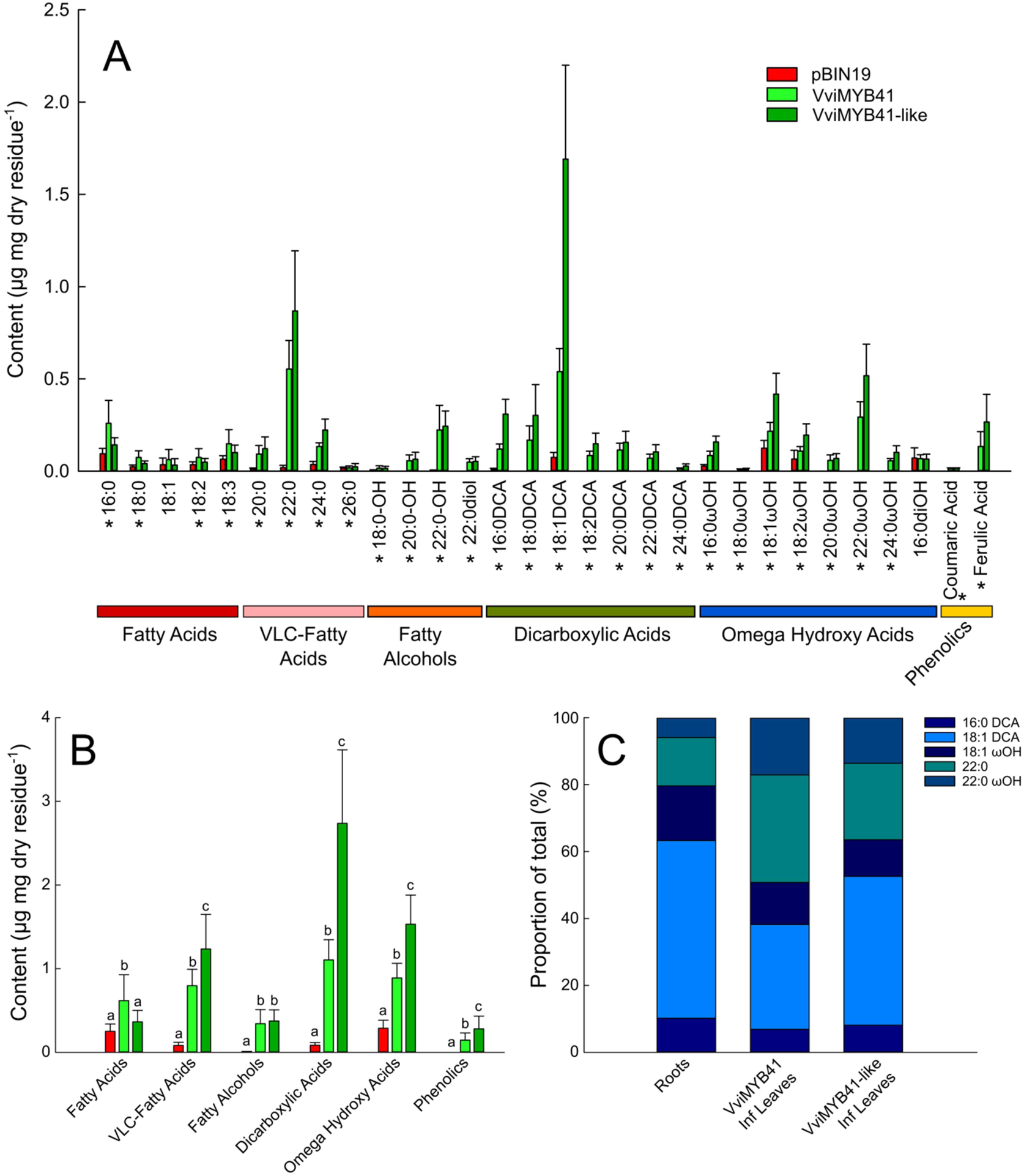
Transient Expression of VviMYB41 and VviMYB41-like in *Nicotiana benthamiana* leaves. Suberin monomers from solvent-extracted leaves were released by transmethylation, silylated, separated by gas chromatography and quantified using internal standards. A, Monomer composition and content of control (pBIN19, red bars), VviMYB41 (light green bars), and VviMYB41-like (dark green bars) agroinfiltrated leaves. B, Acyl-chain content of control pBIN19, VviMYB41, and VviMYB41-like agroinfiltrated leaves. C, Relative proportions of the five major monomers in *Nicotiana benthamiana* well-watered primary roots, and VviMYB41 and VviMYB4-1like agroinfiltrated leaves. Error bars represent ± standard error, in A stars designate statistically significant ANOVAs (P<0.05) and in B different letters represent statistically significant differences (P<0.05 TUKEY’s HSD; n=15 pBIN19, n=16 VviMYB41, and n=8 VviMYB41-like).

## DISCUSSION

In the current study, we quantified changes in suberin composition during grapevine fine root development and in response to water deficit, and paired these analyses with a transcriptomic study of suberin biosynthesis- and deposition-related genes. Generally, grapevine suberin composition did not differ between primary and lateral roots, and was similar to that of other species thus far described. Under water deficit there was a global upregulation of suberin biosynthesis which resulted in a higher content of suberin specific monomers, but without changes in the relative abundances of these molecules, and this upregulation took place across all the developmental stages of fine roots. These changes corresponded to the upregulation of numerous suberin biosynthesis- and deposition-related genes which included two orthologs of the previously characterized AtMYB41 transcriptional regulator (Kosma et al., 2014). Functional validation in *Nicotiana benthamiana* confirmed that these grapevine MYB41 orthologs were able to globally upregulate suberin biosynthesis and deposition. This study provides a detailed characterization of the developmental and water deficit induced suberization of grapevine fine roots and identifies important orthologs responsible for suberin biosynthesis, deposition, and its regulation under drought in grape.

### Global Composition and Similarities with other Species

The global composition of suberin in grape fine roots is dominated by the four major suberin monomers that have been described as major components of suberin in other species (16:0-DCA, 18:1-DCA, 18:1-ωOH and 22:0-ωOH). 18:1-DCA and 18:1-ωOH represent, by far, the two major aliphatics present in potato suberin periderm (Serra et al., 2009) while 22:0-ωOH represent the second major monomer (after 9,10-epoxy-octadecane-1,18-dioic acid) of cork oak suberin (Cordeiro et al., 1998). Similarly, Arabidopsis root suberin is dominated by 18:1-DCA, 18:1-ωOH, 22:0-ωOH and 22:0 (Höfer et al., 2008). 16:0-DCA has been reported in these 3 species, but grape fine roots differed in that they contained high amounts of 16:0-DCA, which represented in about 20 % of the total at each developmental stage. In addition, grape root suberin contained high levels of diols, which have been detected in the Arabidopsis seed coat suberin, but not in Arabidopsis root suberin (Li-Beisson et al., 2013). The physiological impact of these differences is currently unknown, but would be of interest for future studies.

### The Role of Suberin as a Barrier during Root Development and in Response to Water Stress

Suberin is well characterized regarding its role as a barrier to water, solutes, gases, and pathogens in a variety of tissues (Franke and Schreiber, 2007; Barberon et al., 2016). In roots, suberin serves as a barrier in the exodermis, endodermis, and periderm (Enstone et al., 2002). In the current study, the root tip contained very few suberin-specific aliphatics, but the level of suberin-specific aliphatics increased along the root length in agreement with previous qualitative, microscopy based studies of grapevine fine roots (Gambetta et al., 2013). Gambetta et al. (2013) demonstrated that the suberization patterns in grapevine fine roots correspond to large differences in root hydraulic conductivity (Lp_root_); more developed root portions had 10-fold lower Lp_root_. To date, this is the first study to carry out a detailed comparison of suberin composition across multiple stages of development in perennial fine roots. However, comparisons in Arabidopsis both along the primary root, or between taproots and younger roots, showed similar differences where suberin aliphatics increased in older root portions (Höfer et al., 2008; Kosma et al., 2012; Delude et al., 2016). In the current study, the relative contribution of specific suberin monomers did not change along the root length. However, the level of the suberin-specific aliphatics did greatly increase along the root length. Interestingly, the amount of hydroxycinnamates, which have been implicated in tertiary suberin structure and sealing properties (Ranathunge et al., 2011; Vishwanath et al., 2015) also increased in the older segments of the root.

Under water deficit root suberization has been shown to increase in grape (Barrios-Masias et al., 2015) and other species (Enstone et al., 2002), and is thought to contribute to the observed decreases in Lp_root_. The same was true in the current study where there was a global increase in suberin-specific aliphatics in response to water deficit in all root portions which was proportional to the level of water deficit, and this increase in suberin corresponded to decreases in Lp_root_. Like the increases in suberin during development, the increases in suberin under water stress were global and the relative abundances of specific monomers did not significantly change, suggesting a global transcriptional regulation. Several other physiological changes in fine roots are thought to contribute to decreases in Lp_root_ including root lacunae formation (Cuneo et al., 2016), root shrinkage, and decreases in the expression and activity of aquaporins (Gambetta et al., 2017).

It should be pointed out that increased suberization of fine roots can result from both increases in the level of suberin itself, but also changes in the proportion of the fine root that is suberized (Barrios-Masias et al., 2015). In our study, each root portion used for suberin composition analysis was confirmed by microscopy and not simply chosen as a function of the distance from the root tip. This strategy circumvented this potential bias and allowed us to compare root portions of similar developmental stages under various levels of water deficit. However, for RNAseq analyses the first 5cm of fine roots were taken, which corresponded to different developmental zones. This is a limitation of the current study and we cannot make detailed conclusions regarding which specific developmental stages contribute to the observed changes in expression for the genes selected.

### Orthologous Gene Candidates for Suberin Lamellae Formation in Grapevine

We analyzed a selection of orthologous grapevine genes involved in suberin biosynthesis and deposition guided by previously characterized orthologs in other species (Vishwanath et al., 2015). Like most gene families in grape and other plant species, each family contained many homologous genes (likely arising from duplication events), in agreement with the idea that homologous genes within a family undergo some sub-functionalization, with particular homologs being expressed in specific tissues, at specific times, or in responses to specific environmental cues (Falginella et al., 2010; Vannozzi et al., 2012; Wong et al., 2016). Sub-functionalization was also evident in the current study where of the 11 gene families we analyzed all had one or multiple specific family members that were expressed in grapevine fine roots.

Many gene candidates that were upregulated under water deficit belong to gene families involved in the biosynthesis (e.g. KCS, CYP86, LACS, etc.), deposition (e.g. ABCG), and regulation (e.g. MYB41) of suberin. This corresponded to a global increase in suberin aliphatics under water deficit. The significant upregulation of the *Vitis* orthologs of the Arabidopsis *AtMYB41*, along with their functional validation in *N. benthamiana*, suggests that these function as global regulators of suberin biosynthesis in grapevine as well (Kosma et al., 2014; Lashbrooke et al., 2016; Legay et al., 2016). The putative grape MYB107 (VIT_16s0039g01710) ortholog identified in the multispecies meta-analyses of Lashbrooke et al. (2016) was expressed in fine roots in this study, but did not change significantly in response to water deficit. However, of the three suberin biosynthesis genes Lashbrooke et al. (2016) utilized as bait in their multispecies co-expression analyses (in grape, GPAT5; VIT_13s0019g01990, FACT; VIT_17s0000g00950, and CYP86B1; VIT_01s0011g02060), both the GPAT5 and FACT orthologues were significantly upregulated under water deficit in this study (the CYP86 ortholog was upregulated as well, but not significantly) confirming the robustness of their analyses. Taken together these results suggest that the biosynthesis and regulation of suberin is well conserved across plant species and that the orthologs identified here play important roles in regulating suberin biosynthesis and deposition in grape, and likely other woody perennials under drought.

### Multiple Transcription Factors Fine Tuning Suberin Deposition

Suberization is a highly dynamic process that is tightly regulated spatiotemporally during development and in response to changes in environment. In roots, this allows for the fine-tuning of suberization as roots adapt to different water and nutrient availabilities. Part of this regulation is hormonal and is mediated through numerous different transcription factors. The induction of suberization by abscisic acid was reported decades ago (Cottle and Kolattukudy, 1982), while its repression by ethylene was recently suggested (Barberon et al., 2016). The direct activation of several suberin-biosynthesis genes by MYB transcription factors has been reported for QsMYB1, AtMYB39, AtMYB92, and AtMYB107 (Gou et al., 2017; Capote et al., 2018; Cohen et al., 2020; To et al., 2020). Recently in kiwifruits, Wei and coworkers showed that abscissic acid induced the expression of *AchnMYB41* which upregulated downstream suberin-related genes (Wei et al., 2020a; Wei et al., 2020b). In Arabidopsis, 6 homologous MYBs that cluster together (MYB9, 39, 53, 92, 93 and 107; subgroups S10 and S24 in Dubos et al., 2010) may regulate suberin deposition under normal conditions, while those from a related sub-group that includes AtMYB41 may activate suberin synthesis under conditions of abiotic stress (Kosma et al., 2014). Interestingly, individual MYBs appear to differently affect specific organs in Arabidopsis. For example, AtMYB107 specifically affects seed coat suberin while AtMYB39 primarily affects root suberin (Gou et al., 2017; Cohen et al., 2020). In addition to MYBs, NAC transcription factors were shown to activate or repress suberization (Verdaguer et al., 2016; Mahmood et al., 2019; Soler et al., 2020), suggesting that multiple transcription factor families collectively contribute to the control the suberization process allowing for the deposition of suberin in specific cell types and in response to specific environmental factors. In our study other MYB transcription factor orthologs of *AtMYB9*, *AtMYB39*, *AtMYB92*, *AtMYB107*, and *MdMYB93* (Lashbrooke et al., 2016; Legay et al., 2016; Gou et al., 2017; Cohen et al., 2020; To et al., 2020) were also expressed in grapevine fine roots, and could therefore be of interest for future study.

Previous transcriptomic analyses identifying the putative downstream targets of some MYB transcription factors demonstrate that they do not solely control aliphatic suberin biosynthesis and deposition. Additionally, these transcription factors also regulate the expression of several genes involved in the phenylpropanoid and cell wall metabolism pathways, suggesting they have a global role in the formation of suberin lamellae (Lashbrooke et al., 2016; Legay et al., 2017; Capote et al., 2018). Their role as master regulators is further supported by the recent discovery that AtMYB92 upregulated the expression of several glycolytic and fatty acid biosynthetic genes, i.e. genes encoding enzymes responsible for producing early precursors of suberin aliphatic monomers (To et al., 2020). Several transcriptomic studies also reported that these suberin-related transcription factors affect each other’s expression (Legay et al., 2017; Cohen et al., 2020). Further studies will therefore be needed to elucidate their hierarchical relevance and potential interactions.

## CONCLUSION

This study provides a first analysis of the aliphatic composition of suberin along grapevine fine roots. It also extends our knowledge of the regulation of suberized root structures in woody perennials and their regulation under water deficit. Our results demonstrate that increases in suberin deposition according to the developmental stages (along the root length) and in response to water deficit appear to be global, involving increases of all aliphatic monomers. In grape, two MYB41 orthologs contribute to the regulation of suberin biosynthesis and deposition in response to drought, suggesting that in some species duplication events can lead to the involvement of multiple MYB41 orthologs. Future studies could focus on more detailed expression analyses, ideally in specific tissues, developmental stages, and in response to other stresses.

## MATERIALS AND METHODS

### Plant Material and Growing Conditions

The commonly used commercial grapevine rootstock RGM (Riparia Gloire de Montpellier, *Vitis riparia*) was used in this study. One-year old dormant grapevine cuttings were purchased and stored in a cold chamber (4°C) until the time of utilization. Before plantation, cuttings were rehydrated for 24 hours at 25°C. Following rehydration grapevines were planted in cylindrical rhizotrons (height 40 cm × diameter 14 cm) with 100 % sand (allowing harvest of the roots without damage) and only one bud at the top node was kept for shoot growth. The plants were grown in a greenhouse at INRA-Aquitaine Villenave d’Ornon, France. Plants were watered until capacity directly after plantation and maintained at capacity via automatic irrigation system with standard nutrient solution. The composition of the nutrient solution was: 2.5 mM KNO_3_, 0.25 mM MgSO_4_-7H_2_O, 0.62 mM NH_4_NO_3_, 1 mM NH_4_H_2_PO_4_, 9.1 mM MnCl_2_-4H_2_O, 46.3 mMH_3_BO_3_, 2.4 mM ZnSO_4_-H_2_O, 0.5 mM CuSO_4_ and 0.013 mM (NH4)6Mo_7_O_24_-4H_2_O (Tandonnet et al., 2009). Iron was supplied as 8.5 mg/L Sequestrène 138 (EDDHA 5.9% Fe) and the final pH of the nutrient solution was 6.0 (Tandonnet et al., 2009).

After an approximate month long establishment period the grapevines were ~1m in height and 10-20 nodes (Supplemental Fig. S5) and were randomly assigned to two water treatments: well-watered and water-stressed. Plants under well-watered conditions were maintained at capacity as during the establishment period and were referred to as control, and plants under water-stressed conditions did not receive any water supply during the period of treatment and were referred to as water-stressed. There were approximately 30 plants in each cohort. Water-stressed plants were binned into two different categories of stress, Medium and High, defined by their predawn water potential (Ψ_predawn_; measurements described below). Medium stress corresponded to Ψ_predawn_ > −1MPa, and High to Ψ_predawn_ < −1MPa. The duration of the dry-down was approximately one week to reach medium stress, and two weeks to reach high stress. Experiments were carried out in both the 2015 and 2016 growing seasons. In 2015, one experiment was used for the suberin polyester comparison of primary and lateral roots (control plants only). In 2016, separate experiments were conducted; one from which physiological measurements of water status, gas exchange, and root hydraulic conductivity were recorded and another from which samples for global lipid analysis of root segments and RNAseq analyses were taken (i.e. global lipid and RNAseq analyses were carried out on different roots respectively, but from the same plants). The water stress treatments were no different between these two experiments (Supplemental Fig. S6).

### Water Status, Gas Exchange, and Root Hydraulic Conductivity

During the experiments water status and gas exchange was monitored periodically on all plants to follow the progression of water deficit. For Ψ_predawn_ measurements a mature leaf from the middle part of the stem was sampled before sunrise. Ψ_predawn_ was determined using a Scholander pressure bomb (Model 1000, PMS Instrument, Albany, OR, and SAM Precis, Gradignan, France). Stomatal conductance and assimilation measurements were conducted using an infrared gas analyzer (GFS-3000, Heinz Walz GmbH, Effeltrich, Germany) using ambient conditions. Measurements were taken only on mature leaves between 8:00 and 11:00AM. Reported Ψ_predawn_, stomatal conductance, assimilation, and root hydraulic conductivity values (Fig. 3) are those recorded on the day that the plants were harvested for analyses (microscopy, suberin analyses, RNAseq).

Hydraulic conductivity of individual roots (Lp_root_) was measured only at the end of the experiment as these measurements are destructive. Lp_root_ was determined using an osmotic pressure gradient with a meniscus tracking method (Gambetta et al., 2013; Knipfer et al., 2015). Root sampling took place between 10h00 and 12h00. Growing medium around one targeted root was carefully removed, and the root was maintained intact. Individual fine roots were cut off with a razor blade under water, choosing roots that were as long as possible while avoiding lateral roots, and measurements were made immediately to avoid artefactual decreases in the measured Lp_root_. Roots were glued into a 500-mm-diameter glass capillary via a home-made adaptor. The capillary was filled with deionized water (diH_2_O) and the water-air interface was observed with a webcam connected to a laptop using YAWCAM (Version 0.5.0). Roots were submerged in a series of aerated sucrose solutions (at least 4 concentrations ranging from 0-0.7MPa) and images of the meniscus were taken every 30 seconds. ImageJ (1.51a, Wayne Rasband) was used to calculate the movement of the meniscus and then the flow rate was obtained. The relationship between flow rate and osmotic pressure was plotted and the slope of the best fit linear regression was the Lp_root_. The length and diameter of each root were measured in order to estimate the root surface area. Lp_root_ was normalized by root surface area.

### Epifluorescence Microscopy

Fresh roots were sampled and kept in 70% ethanol at 4°C for further observations of their anatomical structure. A berberine-aniline blue fluorescent staining method was used to stain root sections (Brundrett et al., 1988). Root cross-sections were taken at different locations along the root. Root segments were fixed in 6 % low gelling temperature agarose and cut into 50 µm thick pieces with a vibrant Microtome with razor blade (Microm 650V). After the staining procedure, root sections were mounted on a slide and observed with an epifluorescence microscope Zeiss Axiophot equipped with the Amira software.

### Root Suberin and Global Lipid Analysis

The roots from the plants were carefully separated from the sand, washed with distilled water, dried with paper towels, and stored at −80°C until use. For suberin polyester analysis, a root section corresponding to 5-20cm from the root tip was used. With a pair of scissors, all lateral roots were separated from the primary root and pooled while the primary roots were cut in 1 to 2 cm segments. Both samples were subsequently delipidated and their suberin composition and content analyzed using the solvent-extraction method as described in Delude et al. (2017). For global lipid analysis of root segments, very thin sections of the primary roots were taken from 0-25cm from the root tip, sliced with a razor blade, and directly observed under a Leica MZ16F Stereomicroscope in order to determine the different developmental stages of the primary root; Primary Xylem, Early Secondary Growth or Late Secondary Growth with periderm (~2.5, 12.5, and 26 cm from the root tip, respectively). About 1 cm sections corresponding to the different stages were collected and subjected to global lipid analysis using the non-extraction method as described in Delude et al. (2017).

### Transient Expression in *Nicotiana benthamiana*

PCR amplification of *VviMYB41* and *VviMYB41-like* ORFs was achieved using Q5 Polymerase, cDNA matrix and primers listed in Supplementary Table S1. PCR products were subcloned were cloned into pDONR221 ENTRY vector by the Gateway recombinational cloning technology using the attB and attP (BP) recombination sites, and verified by sequencing. Selected clones were then transferred in the overexpression vector pK7W2G2D (Karimi et al., 2002) by LR cloning.

*Agrobacterium tumefaciens* was transformed using a standard electroporation protocol and transformants were selected at 30°C on solid Luria Broth (LB) medium supplemented with antibiotics (gentamycin 25μg/mL with spectinomycin 100μg/mL or kanamycin 50μg/mL). Agrobacterium colonies were inoculated into 2 mL LB supplemented with antibiotics and grown at 30°C overnight. Next morning, the optical density (OD) was measured, and transformation cultures were launched in 5-mL LB by adjusting the initial OD_600_ to 0.1. Cultures were grown for about 4-6 hours to an optimal OD_600_ of 0.6-0.8. Cells were sedimented by centrifugation at 7000 rpm for 5 minutes, the supernatant was discarded, and cells were re-suspended in 5mL sterilized H_2_O. To avoid toxicity, the final bacterial OD600 of the infiltration medium was 0.4. The vector containing the p19 protein to minimize plant post transcriptional gene silencing (PTGS) was used for all experiments (Voinnet et al., 2003).

*N. benthamiana* was agroinfiltrated on the abaxial side of the leaves using a 1-ml plastic syringe without a needle as described by Voinnet et al. (2003). Plants leaves were harvested 5 days later, and immediately inubated in isopropanol for 30min at 85°C. Delipidation and suberin composition and content analysis was performed using the solvent-extraction method as described in Delude et al. (2017).

### mRNA Extraction and RNA Sequencing

Root tips of 5 cm were harvested and frozen immediately in liquid nitrogen and kept in −80°C refrigerator until the time of analysis. Frozen samples were ground in liquid nitrogen into powder for RNA extraction. Total mRNA was extracted after Reid et al. (2006) and genomic DNA contamination was removed with the Turbo DNA-free kit (Life technologies, according to manufacturer’s instructions).

RNA-seq was performed using a HiSeq3000 Illumina sequencing platform with paired-end 150 bp (PE150) sequencing strategy. Three replications were set at each water stress level for the total RNA sequencing and RNA-seq analysis. FastQC was used to check the quality of raw reads generated by Illumina. To obtain high-quality clean reads, the raw reads were first trimmed by removing adaptor sequences using the Cutadapt software. Low-quality reads containing more than 20% bases with a Q-value <10 were discarded. The clean reads were then mapped to the 12X.2 version of Pinot Noir derived PN40024 reference genome sequence. Raw counts were determined using Tophat2 and SAMToolsv (Li et al., 2009), generating an overall counts matrix. Only reads with a perfect match or one mismatch were further analyzed and annotated based on this reference genome. Mapped reads per gene were counted with HTSeq‐count (www-huber.embl.de/users/anders/HTSeq/; Anders et al., 2015). Then, the R package edgeR (Robinson et al., 2010) was used to identify differentially expressed genes using a stringent threshold: absolute value of Log Fold Change (LFC) >1 and False Discovery Rate (FDR) <0.01.

### Dendrogram Construction

The grapevine suberin-related gene family sequences were retrieved from the OrcAE 12x grapevine annotation V2 (http://bioinformatics.psb.ugent.be/orcae/) through a combination of keyword and BLAST searches (using default parameters) based off of the family described in (Vishwanath et al., 2015). Arabidopsis sequences were retrieved from TAIR (https://www.arabidopsis.org/). Multiple sequence alignments and dendrogram constructions were carried out with Phylogeny.fr (Dereeper et al., 2008). Each family was split into sub-families for alignments in order to avoid artifacts caused by aligning large groups (Boyce et al., 2015). Sequences were aligned with MUSCLE (v3.8.31) using the highest accuracy default settings. After alignment gaps and/or poorly aligned regions were removed using Gblocks (v0.91b) using the following parameters: minimum length of a block after gap cleaning = 5, no gap positions were allowed in the final alignment, all segments with contiguous nonconserved positions bigger than 8 were rejected, minimum number of sequences for a flank position = 55%. Dendrograms were reconstructed using the maximum likelihood method implemented in the PhyML program (v3.1/3.0 aLRT) using default settings. Reliability for internal branch was assessed using the bootstrapping method (100 bootstrap replicates). Dendrograms were drawn with TreeDyn (v198.3).

### Statistical Analysis

Treatment effects were evaluated using a one-way analysis of variance (one-way ANOVA, p < 0.05, Tukey’s HSD test). All ANOVAs were carried out in R version 3.3.1 (2016-06-21) (R Core Team) and all graphs were made with SigmaPlot (Version 11.0, Systat Software).

## ACKNOWLEDGMENTS

The authors would like to thank the CGFB Bordeaux Metabolome Facility – MetaboHUB (ANR-11-INBS-0010) where all suberin GC-based analyses were performed. The authors would also like to thank all the staff at EGFV who aided with production and care of the plant material used in this study.

